# Aromatic ring flips in differently packed ubiquitin protein crystals from MAS NMR and MD

**DOI:** 10.1101/2022.07.07.499110

**Authors:** Diego F. Gauto, Olga O. Lebedenko, Lea Marie Becker, Isabel Ayala, Roman Lichtenecker, Nikolai R. Skrynnikov, Paul Schanda

## Abstract

Probing the dynamics of aromatic side chains provides important insights into the behavior of a protein because flips of aromatic rings in a protein’s hydrophobic core report on breathing motion involving a large part of the protein. Inherently invisible to crystallography, aromatic motions have been primarily studied by solution NMR. The question how packing of proteins in crystals affects ring flips has, thus, remained largely unexplored. Here we apply magic-angle spinning NMR, advanced phenylalanine ^1^H-^13^C/^2^H isotope labeling and MD simulation to a protein in three different crystal packing environments to shed light onto possible impact of packing on ring flips. The flips of the two Phe residues in ubiquitin, both surface exposed, appear are remarkably conserved in the different crystal forms, even though the intermolecular packing is quite different: Phe4 flips on a ca. 10-20 ns time scale, and Phe45 is broadened in all crystals, presumably due to μs motion. Our findings suggest that intramolecular influences are more important for ring flips than intermolecular (packing) effects.

## Introduction

Aromatic side chains play important roles in proteins. Often located in their hydrophobic cores, they are key to protein stability. Over-represented in protein-protein and protein-drug interfaces, aromatics play an important role in molecular recognition and binding (1), and are often prominent in the active sites of enzymes. The dynamics of aromatic side chains have been intensely studied research for more than four decades (2–5). An important motivation for this interest is the realisation that aromatic side chains are rather bulky, and their motions, particularly rotations of the aromatic rings, require a significant void volume. Consequently, aromatic ring flips are thought to reveal coordinated movement of surrounding residues. Ring flips of His rings have also been studied in the context of functional mechanisms of enzymes and channels (6, 7). Ring flips of Phe and Tyr, i.e. 180° rotations around the χ_2_ dihedral angle, interconvert two indistinguishable states, and the exchange between these is, therefore, not observable by crystallographic methods. Nuclear magnetic resonance (NMR) spectroscopy can probe such motions in quite some detail, including the time scales of ring flips and the amplitudes and time scales of the ring-axis motions. Solution-state NMR provides insights into ring flips because the two symmetry-equivalent spin pairs (at the two H^ε^-C^ε^ positions, or the two H^δ^-C^δ^ positions, respectively, also denoted *ortho*-CH and *meta*-CH) are exposed to different conformational environments. Observing either two distinct sets of peaks, or a single time-average set of peaks, or possibly line broadening due to exchange, provides evidence for the time scale of flips. A growing arsenal of methods allows for quantification of such exchange processes (8–11). Studying the pressure- or temperature-dependence of such parameters sheds light onto the transition state and created void volume involved in ring flips (12–14). A recent study managed to stabilise a transition state of a ring flip which became then amenable to high-resolution structural investigation (15). Nuclear spin relaxation methods (14, 16) provide another avenue to probe ring dynamics of proteins in solution.

Whether crystal packing has an influence on ring flips is poorly understood, largely because these are invisible to crystallography. Magic-angle spinning (MAS) NMR provides atom-resolved insight into protein assemblies, including crystals. It can, thus, shed light onto the impact of crystal packing on motions. MAS NMR has been used for studying dynamics of aromatic rings (5, 17–19) and protein dynamics more generally (see reviews, e.g. references (20–27)). We have recently applied a selective isotope-labeling strategy combined with sensitive proton-detected MAS NMR pulse sequences to quantitatively probe aromatic ring dynamics over a wide range of time scales (28). The approach uses highly deuterated protein samples, in which ^1^H-^13^C spin pairs are introduced at either the C^ζ^ (*para*-CH), the C^ε^ (*meta*-CH) or the C^δ^ (*ortho*-CH) site. Together with MAS frequencies of 40-50 kHz or above, this strategy leads to sensitive high resolution ^1^H-^13^C correlation spectra. Moreover, given the simplicity of the spin system, with well-isolated ^1^H-^13^C spin pairs, it is straightforward to obtain quantitatively accurate dynamics data without any influence of scalar or dipolar couplings to remote spins. In the solid state, the arsenal of methods for probing dynamics is richer than in solution: in addition to chemical-shift based methods and relaxation measurements (also accessible in solution), MAS NMR (i) allows quantifying dipolar couplings and (ii) provides insights into microsecond-millisecond dynamics from experiments that are sensitive to the MAS frequency and radiofrequency (RF) fields, such as NEar-Rotary resonance Relaxation Dispersion (NERRD) experiments (29, 30) (see below). Measurements of dipolar couplings are very useful to learn about motional amplitudes: motion leads to averaging of dipolar couplings, and the averaged dipolar-coupling tensor reflects the conformational space that the inter-atomic vector samples; the methods we employ here allow to even see anisotropy of the underlying motion, that is caused e.g. by two-site ring flips (28, 31). Additionally, MAS NMR relaxation measurements probe dynamics over a broad range of time scales (21–23). Although MAS NMR relaxation experiments are differently sensitive to different time scales (32, 33), in principle any time scale can be probed. In particular, NERRD experiments allow probing whether motions occur on microseconds (μs) or rather nanosecond (ns) time scales (33).

Here, we use MAS NMR together with highly deuterated samples with specific ^1^H-^13^C spin pairs to investigate phenylalanine ring dynamics in the 8.6 kDa protein ubiquitin, crystallized in three different crystal forms, herein called “MPD-ub”, “cubic-PEG-ub” and “rod-PEG-ub”. These names refer to the crystallisation agent, methyl pentanediol or polyethylene glycol, and the crystal shape. The arrangement of the molecules, in particular with respect to the Phe side chains, is displayed in Fig. 1.

**Fig. 1.**
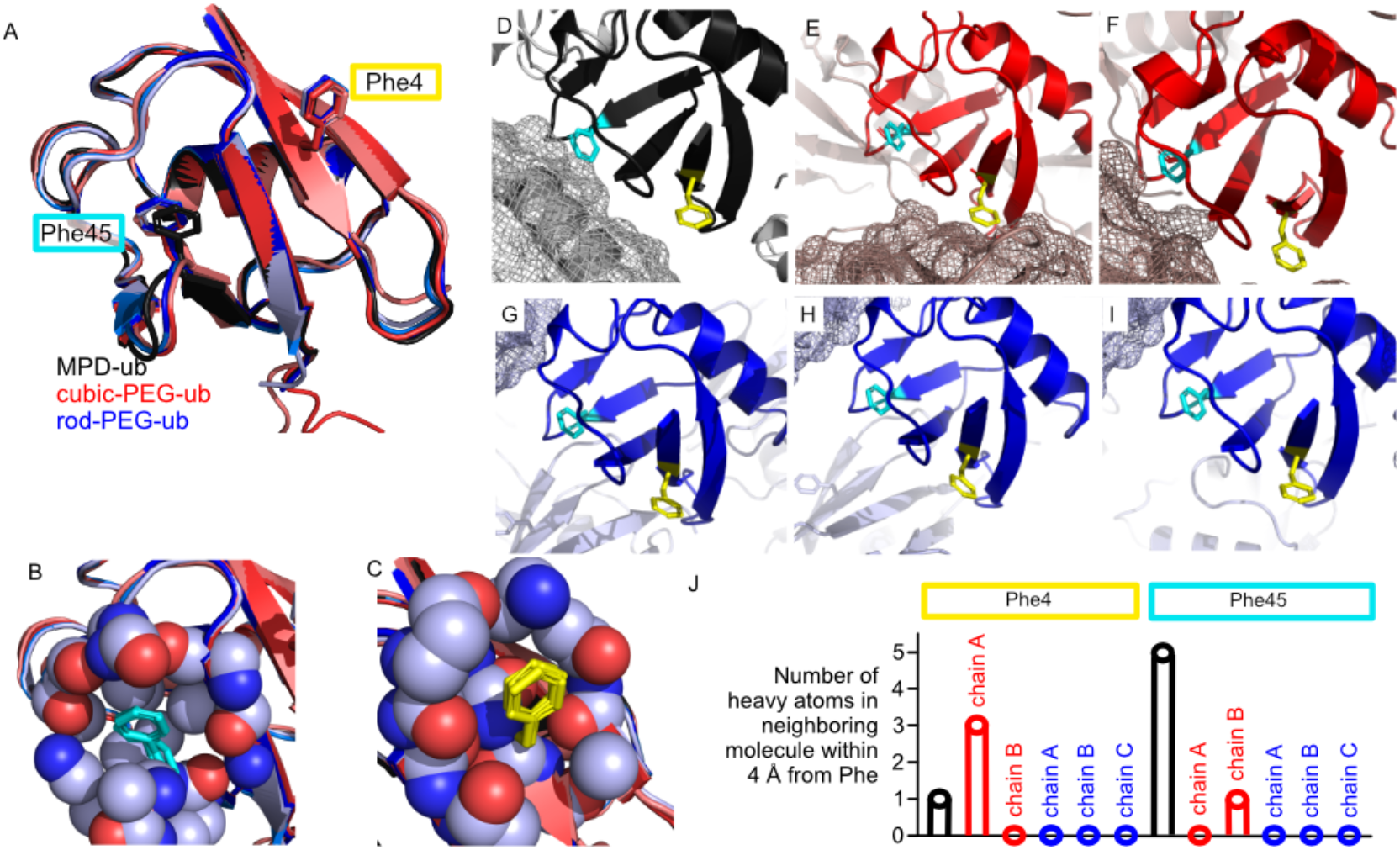
Structures of ubiquitin in three different crystal forms, denoted herein as “MPD-ub” (black), “cubic-PEG-ub” (red) and “rod-PEG-ub” (blue). The three crystal forms correspond to PDB entries 3ONS, 3N30 and 3EHV, respectively. The number of non-equivalent molecules in the unit cell are: 1 (MPD-ub), 2 (cubic-PEG-ub) and 3 (rod-PEG-ub). (A) Overlay of the backbones of the six (1+2+3) chains from the three crystal forms. Panels (B) and (C) zoom onto Phe45 and Phe4, respectively. Spheres denote atoms within 5 Å around the aromatic side chain (shown here for one of the chains of rod-PEG-ub; dark blue: nitrogen, red: oxygen, light blue: carbon). (D-I) Crystal packing for MPD-ub (D), cubic-PEG-ub, chains A (E) and B (F), and rod-PEG-ub, chains A (G), B (H) and C (I). Neighboring molecules in direct contact with the molecule in the center are shown as mesh. (J) The number of heavy atoms located in a neighboring ubiquitin chain within a radius of 4 Å around any atom of Phe4 (left) or Phe45 (right).

These three crystals differ in the number of molecules inside the unit cell, the relative orientation of molecules to each other and the solvent content in the crystal, ranging from 58% water content in cubic-PEG-ub to 49% (MPD-ub) to 40% (rod-PEG-ub). Previous studies of the backbone dynamics of these crystals have revealed that the backbone has similar (sub-μs) motions, but that there is different degree of overall motion: the molecules of the least densely packed cubic-PEG-ub crystal undergo overall rocking motion with an amplitude of several degrees on a μs time scale (29, 34). While both Phe residues are positioned more or less on the protein’s surface, rather than in the hydrophobic core, the packing of the aromatic side chains differ. Using the highly accurate solution-NMR structure of ubiquitin (PDB ID: 1D3Z), we find that the average solvent accessible surface area of Phe4 in this multi-conformer structure is 59 Å^2^ (33%), while for Phe45 it is 42 Å^2^ (23%). However, the degree of solvent exposure changes depending on intermolecular arrangements in the different crystal forms (Fig. 1D-I), which is also reflected in the number of contacts that each Phe side chain makes with the neighboring ubiquitin molecules (Fig. 1J).

MAS NMR combined with MD simulations, presented herein, shed light onto the effects of crystal packing on Phe ring dynamics. The entirety of spectroscopic data suggest that the Phe45 signals are broadened beyond detection in all crystals due to slow (μs) ring flips of Phe45. This finding is backed up by MD simulations. The peak positions of Phe4 are remarkably conserved in the three different crystal forms, despite different buffer composition and pH conditions and intermolecular packing. However, in one crystal form, differences in intermolecular packing between the two chains are reflected by different ^1^H chemical shifts and spin relaxation parameters. Overall, our study reveals that the impact of the crystal packing on aromatic ring dynamics is small compared to the intramolecular determinants of ring flipping. Interestingly, the strong difference of ring-dynamics time scale of Phe4 and Phe45 that we detect from NMR measurements and MD simulations is not directly related to the solvent-accessible surface area, nor the rotameric state, which points to other intramolecular determinants of ring flips.

## Results

### Specifically isotope-labeled Phe samples of three crystal forms of ubiquitin

In order to obtain sensitive and high-resolution proton-detected MAS NMR spectra, combining high levels of sample deuteration with fast magic-angle spinning is an established method (35, 36). For exchangeable sites, in particular the amides of the backbone, perdeuteration followed by back-exchange in ^1^H_2_O buffer is a straightforward method. For detecting side chain atoms, one can either use the “imperfection” of deuteration, i.e. the residual ^1^H content in deuterated samples (37, 38), or specific labeling with precursors in which a chosen type of moiety is protonated almost completely. The latter approach is commonly used for methyl-directed NMR, particularly in solution NMR (39, 40), and also in MAS NMR (31, 41). Introduction of such isolated protons in other sites allows to obtain highest resolution for other side chains; generally, such specific labeling approaches can clearly achieve better line widths than those obtainable from fully protonated samples at the highest available MAS frequencies to date (100 kHz) (28).

We have recombinantly expressed ubiquitin in which all non-exchangeable sites are deuterated, and individual ^1^H-^13^C spin pairs are incorporated at the two H^ε^-C^ε^ positions of Phe residues. We denote this labeling here as u-[^2^H,^15^N],Phe(ε1,ε2)-^1^H,^13^C. (None of the other carbon sites is ^13^C labeled). The incorporation of the specific label was achieved by adding a properly labeled ketoacid precursor molecule (35 mg per liter of culture), displayed in Fig. 2A, to the bacterial culture prior to induction (42), as described in the Methods.

**Fig. 2.**
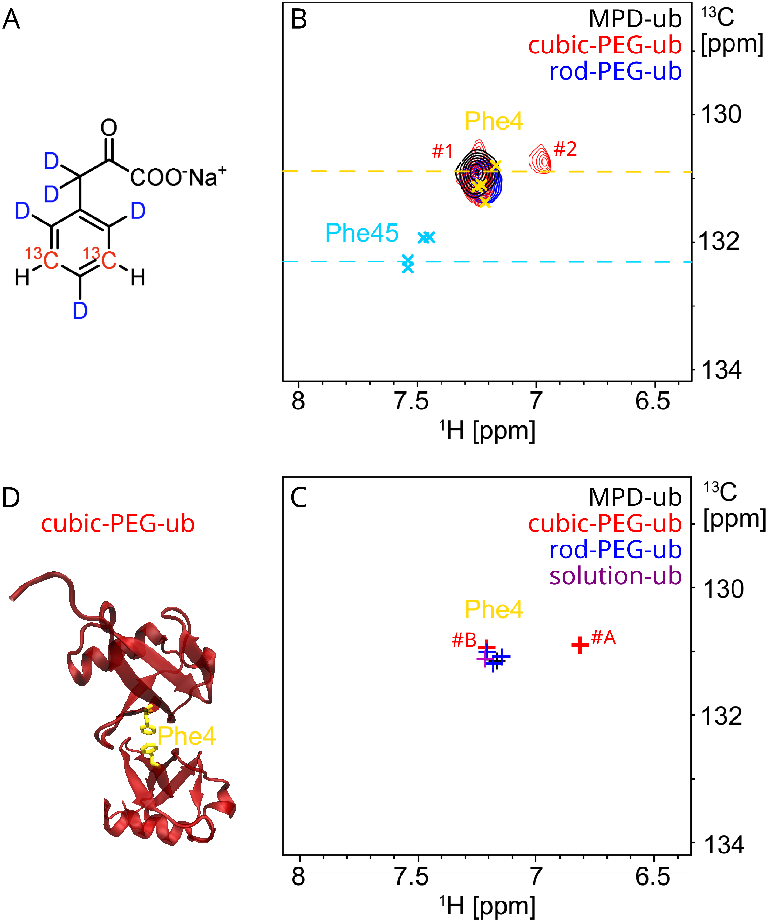
MAS NMR of specifically Phe-labeled ubiquitin. (A) Ketoacid precursor used for the labeling of Phe with two ^1^H^ε^ -^13^C^ε^ spin pairs at the two ε positions. (B) Overlay of the cross-polarization (CP) based ^1^H ^ε^ -^13^C ^ε^ correlation spectra of the three different ubiquitin crystal samples. The spectra were obtained at 50 kHz MAS frequency at a sample temperature of ca. 28 ° C with a pulse sequence based on ^1^H^ε^ -^13^C^ε^ out-and-back cross-polarisation steps and ^1^H detection (28). MPD-ub and rod-PEG-ub spectra feature a single observable peak, while cubic-PEG-ub displays two peaks (labeled #1 and #2 in the plot). The assignments of Phe4 and Phe45 in various solution-state data sets (BMRB 6457, 5387, 16228 and 27356) are indicated by crosses, and the solid-state NMR assignment (only for ^13^C; BMRB 25123) is indicated by a horizontal line, color-coded for Phe4 (gold) and Phe45 (cyan). (C) Chemical-shift predictions for ^1^H^ε^-^13^C^ε^ spin pairs in Phe4 using the program SHIFTX2. Tthe peaks are color-coded as indicated in the legend. Of interest, chain A in cubic-PEG-ub shows a distinctive proton chemical shift, significantly different from that of chain B. This effect can be attributed to ring current shift arising from the stacking of the two Phe4 rings from the proximal chains A in the crystal lattice (illustrated in panel D).

Fig. 2B shows the aromatic region of the ^1^H-^13^C spectra of ubiquitin in the three different crystal forms. Overlaid with these spectra are previous assignments of the ^1^H^ε^-^13^C^ε^ sites in solution (43–46), and of the ^13^C^ε^ in a carbon-detected MAS NMR study (47). Even though these previous data sets have been collected under a diverse set of conditions, including in-cell NMR and a sample in reverse micelles, the reported peak positions are remarkably similar.

### Phe4 rings in all crystal forms undergo sub-millisecond ring flips

In our spectra, only the signal that corresponds to Phe4 is visible, while the one of Phe45 is absent. We discuss the question of why the signal of Phe45 is not observed further below. We observe a single peak for Phe4 in both MPD-ub and rod-PEG-ub spectra. For MPD-ub, in which all chains are identical, this is expected. However, this outcome is less obvious for rod-PEG-ub, where the crystal contains three non-equivalent molecules in the unit cell and many backbone amide sites show three distinct signals (34). Closer inspection of the three non-equivalent Phe4 appearances in the crystal (Fig. 1G–I) shows that the Phe4 side chains do not have any contact to other molecules. Thus, their environment is very similar in the three non-equivalent molecules, which explains the observation of a single signal. In cubic-PEG-ub we observe two signals: one overlaps with the position found in the other crystal forms and in solution (denoted as #1 in Fig. 2), and the other one shifted by ca. 0.25 ppm upfield in proton dimension (#2). These two peaks mirror the differences in the environment of Phe4 in the two non-equivalent molecules in the crystal (Fig. 2E, F, J). We tentatively assign peak 1, which is very close to the solution-NMR position, to chain B, because in chain B Phe4 does not form any intermolecular contacts, and peak 2 to chain A, which is engaged in contacts to a neighboring chain.

To confirm this view, we have performed SHIFTX2 predictions (49) of the ^1^H^ε^-^13^C^ε^ correlations in solution and in the three crystal forms. Chemical-shift prediction programs are often challenged to predict proton frequencies with accuracy, particularly those in side chains, for which fewer data are available. However, the predictions are able to reproduce the effects observed for Phe4 rather well. In particular, the prediction finds a 0.39 ppm upfield shift of the signal corresponding to chain A of cubic-PEG-ub, compared to chain B. This effect is due to intermolecular stacking of Phe4 rings in the two neighboring chains A (Fig. 2D). When the second chain is removed in the SHIFTX2 calculation, the two Phe4 signals land essentially on top of each other. Even though predictions of proton chemical shifts are not very reliable, the effects of ring-currents are well understood (50, 51), and we assume that the effects of intermolecular ring stacking are well recapitulated in the predicted Phe4 shifts of cubic-PEG-ub.

On the other hand, the limited accuracy of the structure-based chemical shift predictors is apparent in the results for Phe45. While SHIFTX2 correctly predicts that Phe45 shifts are similar among the different crystal forms and in solution, their absolute values do not agree very well with the experiment, see Fig. S1.

The observation of a single cross-peak for Phe4, which is labeled at the two e sites, suggests that the isotropic chemical shift of the two positions is averaged by sub-millisecond ring flips. To gain a more direct insight into ring flips, MAS NMR can probe the dipolar-coupling averaging. The ^1^H-^13^C dipolar-coupling tensor is averaged by motions faster than ca. 10-100 μs (Fig. 28 in ref. (22)). For the case of ring flips the tensor anisotropy (often denoted as the “dipolar-coupling strength”) is reduced to theoretically 62.5% (order parameter S=0.625); moreover, the dipolar-coupling tensor, which is uniaxial in the rigid-limit case (i.e. axially symmetric), becomes biaxial. (We use here the term biaxial; this property is also termed tensor asymmetry in the literature, with an asymmetry parameter, η, where η=0 denotes an axially symmetric tensor. As “biaxial” more precisely reflects the shape of the tensor, and in line with the use in other fields of physics, (52) we use the term biaxiality herein.) One can show that the ring flips not only reduce the anisotropy of the dipolar-coupling tensor to S=0.625, but also increase the biaxiality parameter to a value of η= 0.6 (28). An adapted version of the Rotational Echo DOuble Resonance (REDOR) experiment (53) allows determining order parameters and tensor biaxiality parameters (31). Fig. 3A shows the REDOR curves for Phe4 in the three crystal forms. In all cases, the obtained tensor parameters are in agreement with the ring flip model, i.e. the order parameter matches (within error bars) the expected value of 0.625, and the confidence interval of the biaxiality parameter encompasses the expected value of 0.6, see Fig. 3B. In some crystal forms the order parameters and the biaxiality are somewhat lower than in others, possibly due to additional small-amplitude motions, but it is clear that within error bar the ring flips alone can account for the experimental observations.

**Fig. 3.**
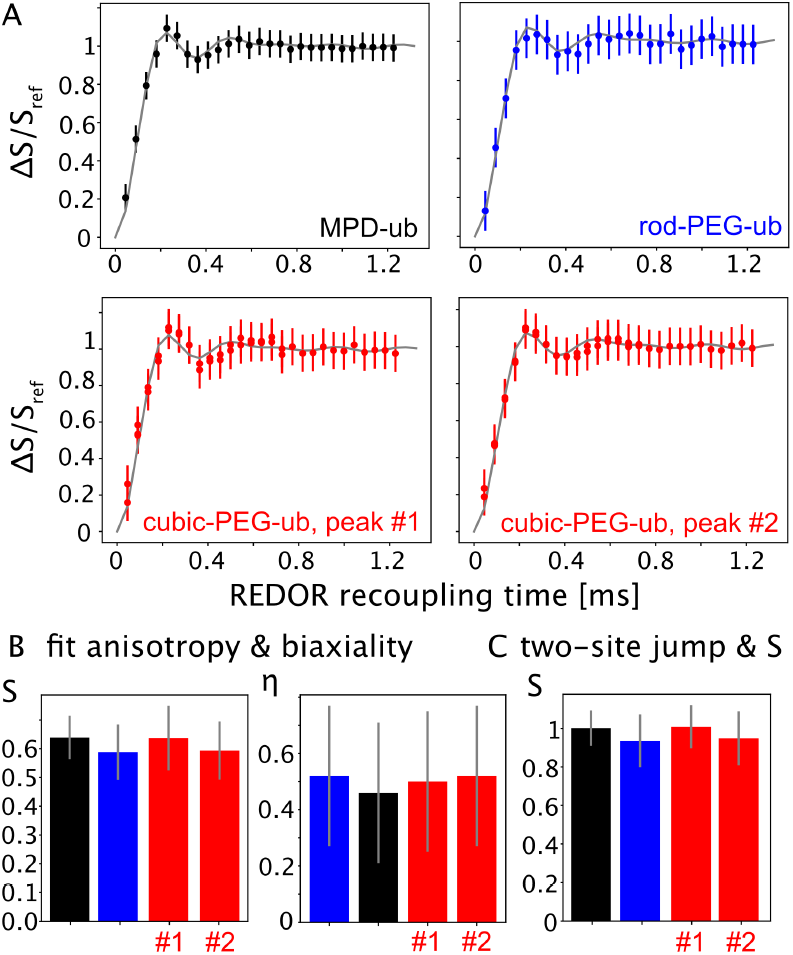
^1^H-^13^C dipolar-coupling tensor measurements for the ε site in Phe4. (A) REDOR recoupling curves for the different crystals/sites. (B) Fitted tensor parameters in a fit that does a grid search for the best order parameter S (i.e. the tensor anisotropy) and tensor biaxiality parameter η. (C) Fitted order parameter from a grid search against a grid of simulations that assume explicit two-site jumps (120°) and a variable tensor anisotropy δ_D_. The resulting best-fit order parameter (calculated, as usual, as S=δ_D,fitted_/δ_D,rigid_) is close to 1, reflecting that the tensor is controlled by the two-site jumps, with only very small additional motional averaging.

### Phe4 ring flips occur on a 10-20 ns time scale in all crystal forms

Spin relaxation rate constants are sensitive to amplitudes and time scales of motion. We have measured the ^13^C longitudinal (R_1_) and rotating-frame 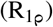 relaxation, as well as ^1^H 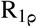, and used them to determine the ring-flip rate constants. To this end, we compared the experimentally measured ^13^C relaxation rate constants (Fig. 4A,B and Tab. S1) to calculated rate constants that result from ring flips (Fig. 4D). In these calculations, we fixed the order parameter of the ^1^H-^13^C moiety to the one expected for ring flips, and varied the corresponding ring-flip correlation time. The calculated relaxation rate constants for a correlation time in the 10-20 ns range match the experimental ones for all crystal forms and for the two sites of cubic-PEG-ub. The correlation times of the ring flips determined from 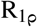 and R_1_ agree very well with each other, supporting that a single dominant motion, ring flips, accounts for the bulk of spin relaxation. We have considered the possibility that additional motions, other than ring flips, contribute to the observed relaxation rate constants. To explore this possibility, we have extended our model by including additional motional modes as found in our MD simulations (see Fig. S2 and Supplementary Note 2). Re-analyzing ^13^C^ε^ *R*_1_ and 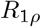 using this extended model shows that additional motions have only minimal influence on the extracted ring-flip rates, see Fig. S3.

**Fig. 4.**
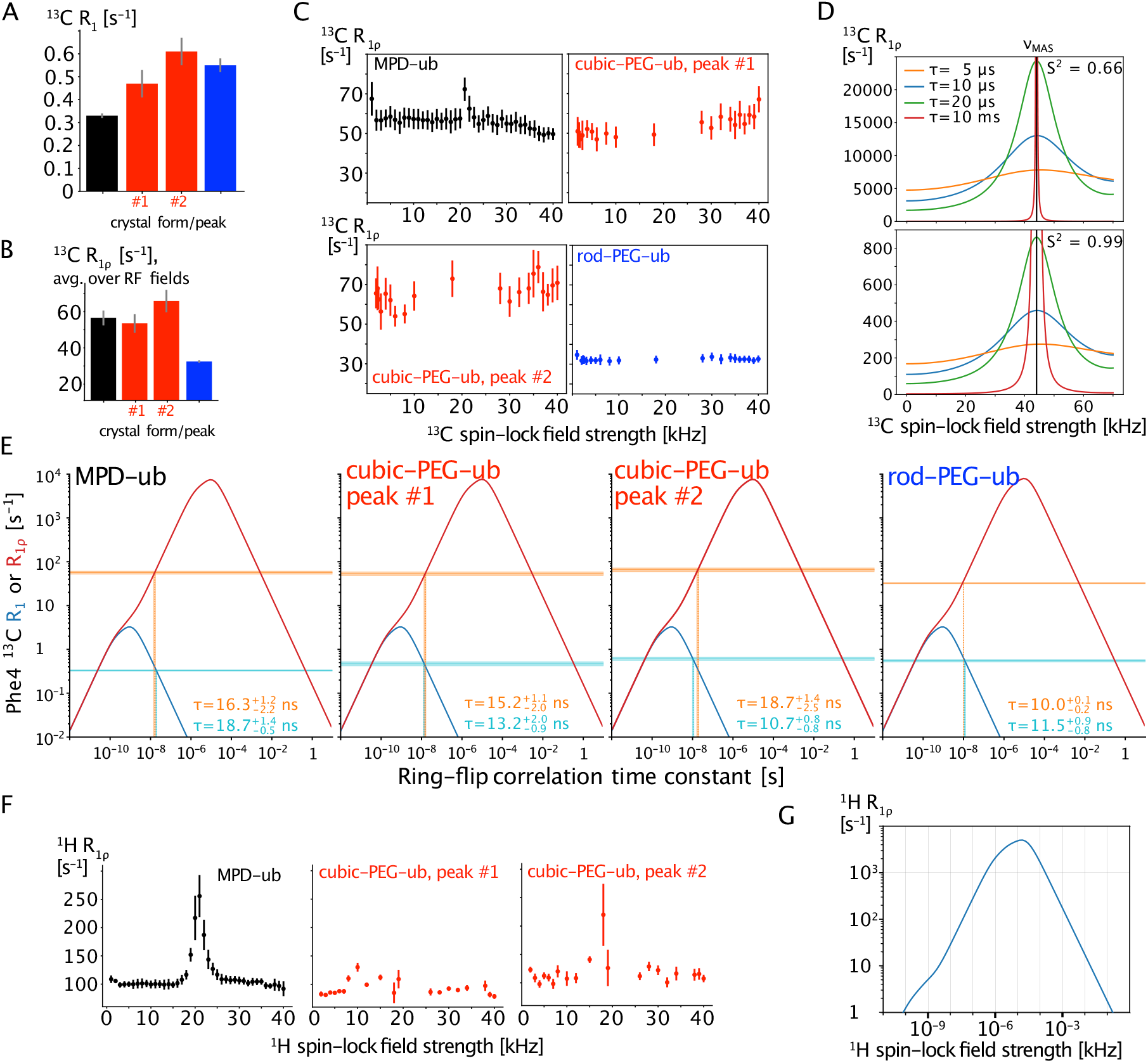
MAS NMR dynamics data for Phe4 signals in the three different crystal forms, sensitive to the amplitude and time scale of motion; all data have been collected at 44.053 kHz MAS frequency. In all cases, the colors black, red and blue refer to data from the three different crystal forms. (A) ^13^C longitudinal (R_1_) and (B) rotating-frame 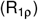 relaxation rate constants. (C) Measurements of ^13^C 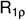 as a function of the spin-lock field strength (relaxation dispersion). The profiles do not show a marked increase when the RF field strength approaches the condition ν_RF_=ν_MAS_ (NEar Rotary resonance Relaxation Dispersion), which one would expect if the flip motion was on the μs time scale. (D) Calculated ^13^C NERRD profiles (see Supplementary Note 1) for different correlation times and an order parameter corresponding to ring flips (S=0.66, upper panel) or smaller amplitude motion (S=0.99, lower panel). The rotary resonance condition is indicated with a black vertical line at 44.053 kHz. (E) Determination of the ring-flip correlation times for Phe4 from ^13^C relaxation rate constants. The Δ -shaped profiles show relaxation rate constants calculated for ring flips (see Supplementary Note 1) as a function of the time scale of these flips; 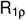; blue: R_1_. Horizontal lines indicate the experimentally measured rate constants and vertical dashed lines show where the experimental data intercept the calculated curve (on the “fast” branch of the curve). (The numerical values of the experimental relaxation rates are summarized in Tab. S1; for 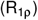, the value at 30 kHz spin lock has been used.) Note the remarkable agreement of the flip time constants from 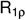 and R_1_. (F) ^1^H 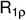 relaxation-dispersion profiles. There is a rise of 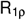 at 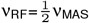 (“HORROR-condition” (48)), which is due to the recoupling of the ^1^H-^1^H dipolar coupling, visible in particular for the MPD-ub crystal data. The width of this feature is limited to only a few kHz around the halved spinning frequency. (G) Calculated ^1^H 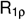 rate constants in the presence of ring flips, taking into account the^1^H CSA and the dipolar coupling to the directly bonded ^13^C and two additional remote ^1^H spins (see Supplementary Note 1 for details).

Even though these data clearly point to nanosecond flips, we also probed whether Phe4 undergoes μs motions, possibly of very small amplitude, using ^13^C 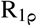 NERRD experiments (29, 30). In these experiments, the 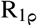 rate constant is probed as a function of the spin-lock RF field strength. In the presence of μs motion, 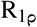 increases sharply when the RF field strength approaches the MAS frequency. A ^1^H 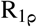 NERRD version has been proposed, too, and, although less straightforward to quantify, is another way to detect μs motions (54, 55). Figs. 4C and F show the ^13^C and ^1^H NERRD data, respectively. These profiles are flat in most cases, suggesting that there is no significant μs motion. The observed rise in the ^1^H NERRD profiles at the condition 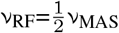, is due to the recoupling of the homonuclear dipolar coupling at the HORROR condition (48); however, it extends over only a narrow range of RF field strengths, much less than what is expected if the motion occurred on a μs time scale (55). It is noteworthy that the experimentally observed ^1^H 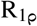 rate constants for Phe4, ca. 50-120 s^−1^, are higher than the expected ones for ring flips occurring on the ca. 10-20 ns time scale (ca. 30-50 s^−1^; Fig. 4G). This suggests that dipolar dephasing (56, 57) appears to be responsible for more than half of the expected decay rate constant.

The calculations illustrated in Fig. 4 also offer a plausible explanation why the signal of Phe45 is unobserved. If ring flips for Phe45 are 1-2 orders of magnitude slower than for Phe4 (MD suggests a factor of ca. 50; see below), then the relaxation time constants of ^1^H and ^13^C at this site are expected to be of the order of 1 ms or less (and additional dipolar dephasing would shorten the ^1^H life time even more). Such fast relaxation would not lead to coherence transfer through the experiment, and broaden signals beyond detection. Hence, Phe45 magnetization would decay rapidly during the pulse sequence and detection, obliterating the spectral signal.

Of note, in cubic-PEG-ub peak #2 (assigned to chain A) has higher ^1^H and ^13^C 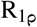 rate constants than peak #1. This is likely the basis why the peak intensity of peak #2 is lower. Given that the ring flip rates are very similar, the origin of this faster relaxation is not entirely clear; a likely reason could be the closer proximity of protons from the neighboring molecule in the crystal, see Fig. 1J.

Lastly, it is noteworthy that the molecules in the cubic-PEG-ub crystals undergo rocking motion within the crystal, while for the more densely packed MPD-ub and rod-PEG-ub crystals this was not found (29, 34). This rocking effect, occurring on a time scale of tens of μs, was detected by non-flat ^15^N NERRD profiles. In this sense, it is interesting that the ^13^C NERRD profiles of cubic-PEG-ub, but not of the other two crystal forms, show a slight (ca. 10 s^−1^) increase toward ν_RF_=ν_MAS_ = 44 kHz. We ascribe this small NERRD effect to the overall rocking motion sensed by the side chain; the effect is quantitatively somewhat smaller than expected from the rocking parameters obtained for the backbone, which might reflect that the side chain is to some degree decoupled from the motion that the backbone senses. It is also possible that the contribution from rocking motion is more difficult to see for these aromatic _1_H-^13^C sites than for backbone amides, because the absolute rate constants are ca. 10-fold larger than those of amide ^15^N.

Overall, NMR data show that in all crystal forms the flips of Phe4 are similar, occurring on a 10-20 ns time scale. It appears that the flips of Phe45 occur on a slower time scale, which prevents its detection by NMR. The MD evidence for that is reviewed in the next section.

### MD simulations provide semi-quantitative insight into ring-flip dynamics

Molecular dynamics simulations provide a useful additional view on ring flips. We have analyzed microsecond-long trajectories of the explicit crystal lattices, as well as of ubiquitin in solution, in order to understand the observed differences in ring-flip rates, in particular between Phe4 and Phe45; the latter is experimentally not observed. Fig. 5 shows the time traces of the χ1 and χ2 dihedral angles of the two Phe sidechains in the different crystal lattices and in solution. As the simulated crystal lattices comprise 24 or 48 molecules, and the simulation extends over 2 μs, the trajectories effectively cover many microseconds.

**Fig. 5.**
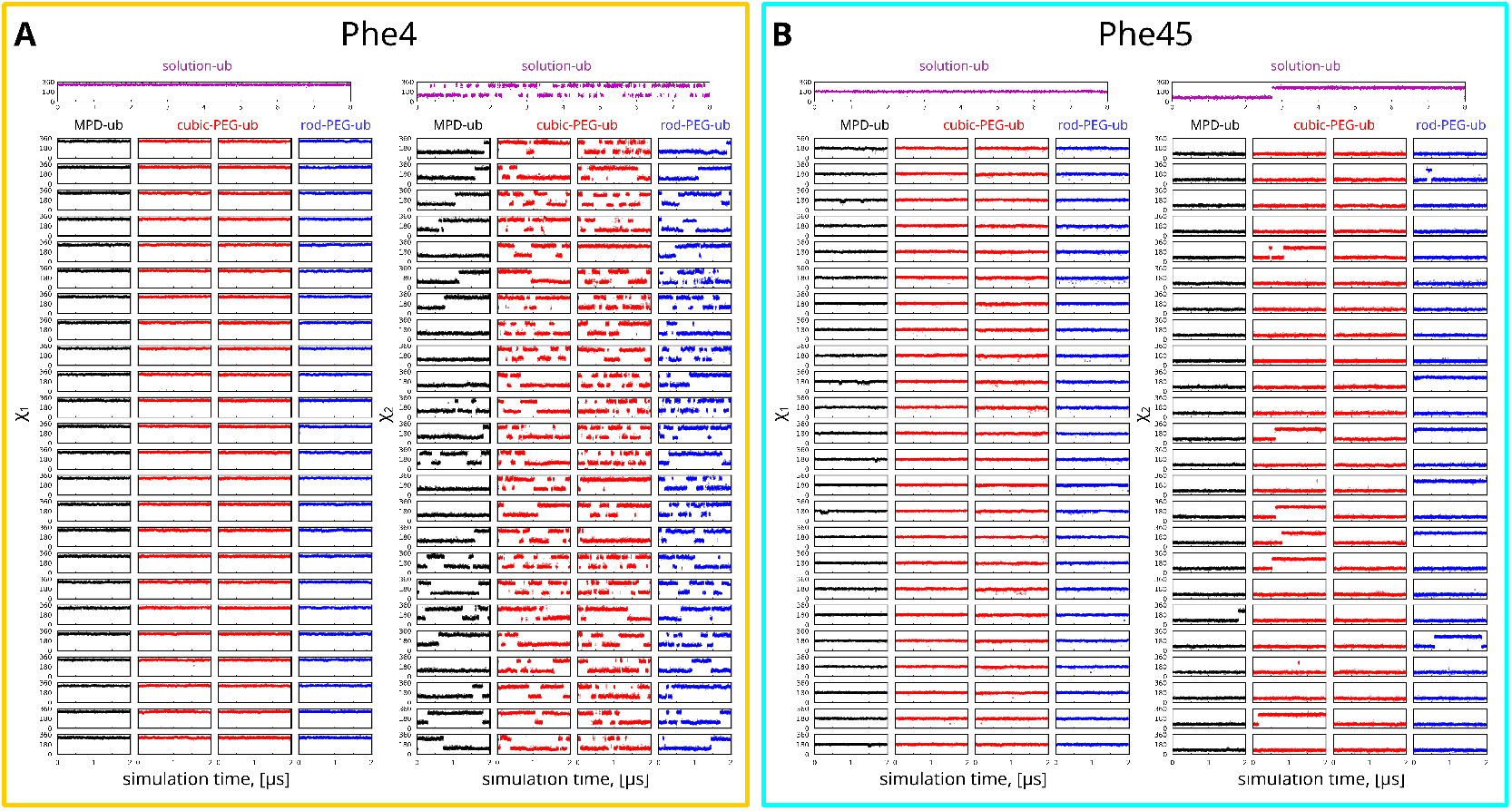
Time traces of side-chain torsional angles χ1 and χ2 for (A) Phe4 and (B) Phe45 from the three simulated ubiquitin crystals and one simulation of ubiquitin in solution. To better visualize rotameric jumps, we use the angle range [0–360°] instead of the conventional choice [-180–180°]. The color coding is the same as in the previous figures: black (MPD-ub), red (cubic-PEG-ub), blue (rod-PEG-ub) and magenta (solution form). The cubic-PEG-ub simulation cell contains 48 ubiquitin molecules equally divided between chain A (molecules 1–24) and chain B (molecules 25–48). The rod-PEG-ub simulation cell contains 24 ubiqutin molecules equally divided between chain A (molecules 1–8), chain B (molecules 9–16) and chain C (molecules 17–24). The details on crystal simulations are provided in the Methods section.

The χ1 angle, which reorients the ring axis C^β^-C^γ^, does not change in any of the simulated systems. For χ2, which represents rotations of the ring (flips and small-amplitude motions within the potential energy wells), the situation is more interesting and more diverse. Phe4 undergoes ring flips in all cases. In cubic-PEG-ub and rod-PEG-ub, these flips occur multiple times along the 2 μs long trajectory. In MPD-ub, which experimentally behaves very similarly, the flips occur less frequently than in the PEG-ub crystals. The aggregate estimate of the characteristic time for Phe4 ring flips using all crystal trajectories and solution trajectory is *τ*_*div*_=204 ns (see Tab. S2 for the flip rate constants of the individual simulations; *τ*_*div*_ is the total aggregated simulation time divided by the number of observed flips). This is an order of magnitude longer than the value of 10-20 ns estimated from the experimental relaxation data. Such difference translates into excess barrier height of ca. 1.5 kcal/mol, which is common even for state-of-the-art force fields (59).

In all crystals as well as in solution, the ring flips of Phe45 are much less frequent than those of Phe4. Specifically, the MD data indicate that there is a 47-fold reduction of the rate constant. While this value is to be considered a rough estimate, due to the insufficient sampling of the flip events in Phe45, the MD data show unambiguously a slow-down compared to Phe4. This kind of slow-down shifts the process into a range that is expected to cause a dramatic broadening of Phe45 signals, preventing its detection in MAS NMR spectra, as further discussed below.

## Discussion

We have shown that a selective isotope labeling scheme with proton-detected MAS NMR provide insights into phenylalanine dynamics in ubiquitin crystals. The resolution in the ^1^H dimension allowed us to detect two distinct environments of Phe4 in the two chains in the asymmetric unit cell. Specifically, stacking of Phe4 rings between two neighboring molecules leads to an upfield proton shift, thus giving rise to a distinct signal for one of the two chains in cubic-PEG-ub. In rod-PEG-ub, which is composed of three non-equivalent chains, only one peak is seen because Phe4 points into the solvent in all chains. Phe45, unobserved in all three crystals, is presumably broadened beyond detection by the slow flips, as suggested by MD simulations. Using the experimentally determined time scale of ca. 10-20 ns for the flips of Phe4 (Fig. 4E) and the MD-derived factor by which Phe45 is slower than Phe4, ca. 50, we estimate that Phe45 ring flips occur on a time scale of about 1 μs.

We have calculated proton transverse relaxation rate 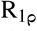 (using a 4-spin system, see Supplementary Note 1 for details), and carbon 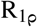 as a function of the time scale of dynamics, to estimate how rapidly the spin magnetization would decay if ring flips occur on such long time scales. These data (Fig. 4E and G) show that the coherence life times under a spin lock are a fraction of millisecond. Such a rapid decay means that the signal is expected to be very broad, and during the coherence transfer steps in an hCH experiment it should die off before detection. In a simple proton single-pulse excitation 1D spectrum, the aromatic signals overlap with the amide signals, making it impossible to detect Phe45. (Deuterating fully all amide sites, i.e. using 100% deuterated buffer, is not possible as even the precipitation agent, MPD, brings in about 20% ^1^H into the solvent.)

It is noteworthy that Phe45 has been detected in one MAS NMR study of MPD-ub, although at lower temperature (273 K) and using carbon detected experiments (47). While we can only speculate about the reasons, a possibility is that Phe45 in MPD-ub has ring flips that are much slower than in other crystals, e.g. in hundreds of μs. This conjecture finds some support in our MD simulations of MPD-ub crystal, where Phe45 shows only one single flip (see Fig. 5B). If so, then lowering the temperature to 273 K may further slow down Phe45 flips, bringing them to millisecond territory (60). That would create the conditions for Phe45 signal to become observable again, cf. Fig. 4E and G. In this connection it should be mentioned that under the conditions of fast MAS used in our study, the lowest temperature that we could achieve was ca. 15-20° C, and we did not observe the Phe45 peak in these trial experiments (not shown).

Of note, the B factors of the δ and ε carbons that sense flips are comparable for Phe4 and Phe45 in each of the crystals (Fig. S4). It is of course not surprising that the strong differences that we find in NMR data are not seen by X-ray crystallography: the difference between Phe4 and Phe45 is the time scale of flips, and crystallography cannot see flips nor their time scale; at cryogenic temperatures they are furthermore expected to be frozen out.

We sought to identify the origin of the large difference in ring-flip rates between Phe4 and Phe45. Given that Phe4 and Phe45 are not buried in the hydrophobic core, but are positioned on the outside of the ubiquitin molecule, a parameter that likely has an influence is the solvent accessible surface area (SASA). We reasoned that SASA should reflect the void (i.e. in this case water-filled) volume available to the phenyl ring. Another factor that might influence the flip rate is the arrangement of phenylalanine side chain relative to the backbone, i.e. its rotameric state. Fig. 6 summarizes structural information and MD data on rotameric states and SASA of the two phenylalanine residues in ubiquitin. The heat maps in this figure show that Phe4 and Phe45 populate two distinct rotameric states, possibly offering an explanation for the observed differences in ring-flip rate constants. To test this proposition, we looked at surface-exposed Phe residues in MD trajectories of several unrelated proteins. In doing so, we found phenylalanines that belong to the same rotameric state as Phe4 but do not experience any flips, as well as other phenylalanines that belong to the same rotameric state as Phe45 but undergo frequent flips (Fig. S5).This observation rules out the possibility that the rotameric states can explain the differences in ring flips. Fig. 6 also shows SASA distributions as obtained from our MD simulations, as well as SASA values from solution and crystal structures (in addition, SASA variations on per-molecule basis are illustrated in Fig. S6).No clear correlation is found between the SASA characteristics and the observed ring-flip rates. Thus, we are led to conclude that the flip rates are likely controlled by a mix of structure/dynamics factors, involving phenylalanine residues and their immediate surroundings, that remain to be fully identified.

**Fig. 6.**
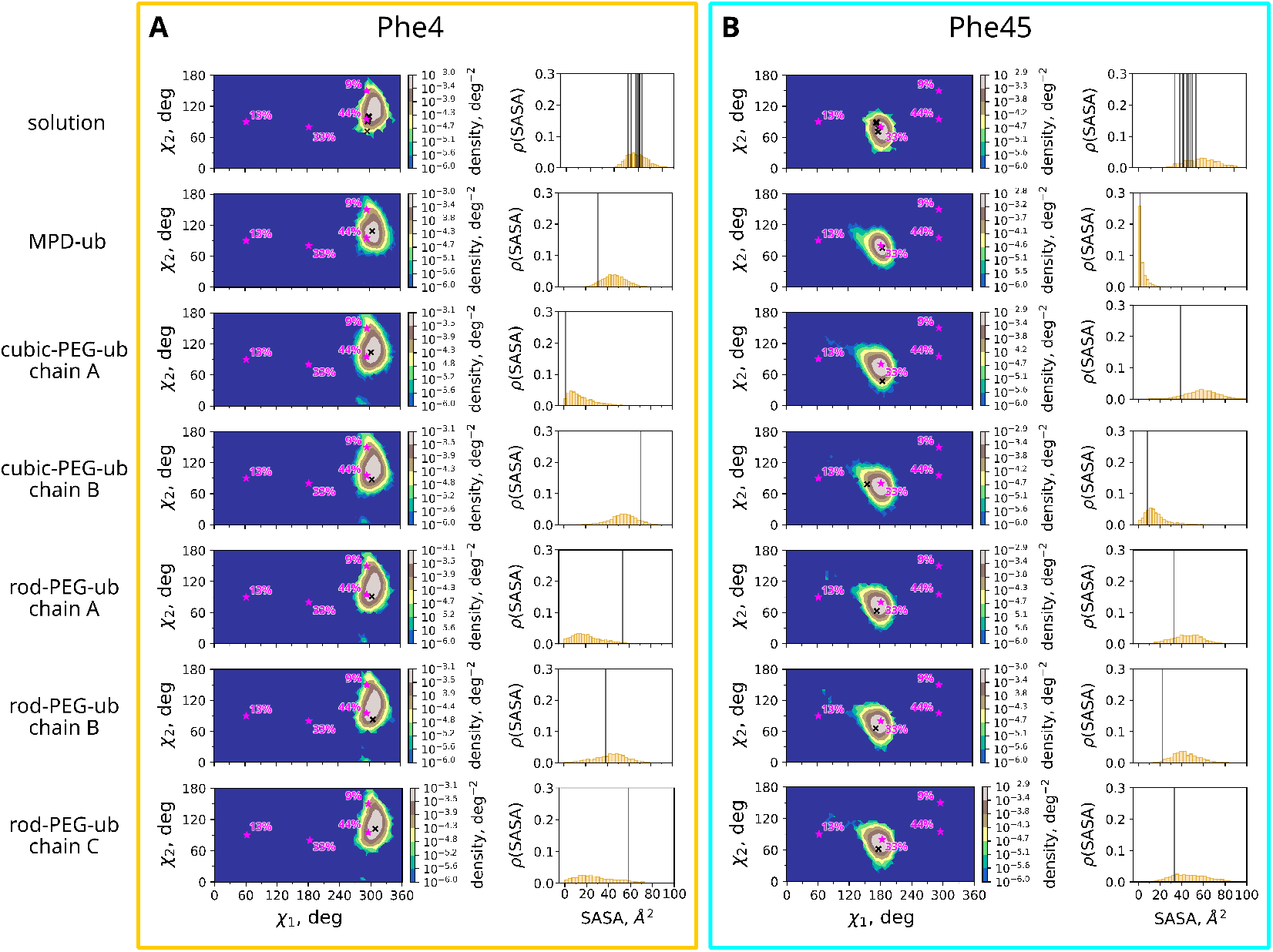
Rotameric states and solvent-accessible surface areas for residues (A) Phe 4 and (B) Phe45 from MD simulations and various structural data. Respective crystal and solution species are identified on the left side of the plot. First column: heat maps showing (χ1,χ2) probability density distribution for Phe4 side chain as obtained from our MD data. Shown with black crosses are the values of χ1, χ2 according to solution NMR (1D3Z, 10 conformers) or crystallographic (3ONS, 3N30, 3EHV) structures. In addition, canonical phenylalanine rotamers according to ref. (58) are indicated with magenta stars along with their respective frequencies of occurrence. The χ1 and χ2 range is [0÷360°], same as in Fig. 5. Second column: histograms showing SASA distributions for Phe4 residue as obtained from our MD data. Also shown are Phe4 SASA values from the solution NMR and crystallographic structures (black vertical lines). Third and fourth columns: the same data for ubiquitin residue Phe45.

Interestingly, while a previous study has found overall rocking motion in cubic-PEG-ub crystal, which leads to strong NERRD effects and overall higher 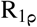 relaxation rates, we do not find significantly different relaxation for the Phe carbons of that particular crystal, and any NERRD effect seen therein is weak, if not absent (Fig. 4C). It is possible that the effects of rocking motion are masked in this case by ring flips.

Overall, our study reveals that for ubiquitin the crystal packing has little effect on ring-flip dynamics: for the observed Phe4, the flip rate constants are nearly the same, and in all crystals Phe45 appears to be broadened beyond detection, which we ascribe to much slower flips. It will be interesting to probe the effects of crystalline packing on ring flips of aromatic residues buried in the hydrophobic protein core, where “breathing” motions are thought to be required for ring flips. We expect that the said breathing motions are largely insensitive to the soft restraining effect of crystal contacts. The methodology we have presented here, which combines selective labeling and a suite of MAS NMR experiments, is well suited to address this question.

## Methods

### Protein expression and purification

Perdeuterated ubiquitin with specific ^1^H-^13^C labels at the phenylalanine e positions was prepared by bacterial overexpression as follows. *Escherichia coli* BL21(DE3) cells were transformed with a pET21b plasmid carrying the human Ubiquitin gene. Transformants were progressively adapted in four stages over 48 h to M9/D_2_O media containing 1 g/L ^15^ND_4_Cl, 2 g/L D-glucose-d_7_ as the sole nitrogen and carbon sources. In the final culture, the bacteria were grown at 37 ° C. When the optical density at 600 nm (OD_600_) reached 0.65, 35 mg of the ketoacid precursor (shown in Fig. 2A) per liter of culture were added. One hour later, while shaking at 37 ° C the OD_600_ reached 0.95, whereupon protein expression was induced by addition of IPTG to a final concentration of 1 mM. Induction was performed for 3 h at 37 °. At the end, the final OD_600_ reading was 2.2.

After induction, the cells were resuspended in 20 mL of 50 mM Tris-HCl pH 8 buffer containing 2 μg/mL leupeptine and 2 μg/mL pepstatine, and lysed by sonication. The lysate was centrifuged for 30 min at 46,000 g using JA25-50 Beckman rotor, then the supernantant was dialyzed against two times 300 mL of 50 mM Tris-HCl pH 8 buffer. After dialysis the sample was centrifuged for 30 min at 46,000 g and loaded on a 40 mL Q-Sepharose column. Ubiquitin was recovered in the flow through fractions, which were subsequently concentrated and injected on a HiLoad 16/60 Superdex 75 column equilibrated with 1 column volume of 50 mM Tris-HCl pH 8 buffer. The protein was dialyzed against Milli-Q Ultra pure water until the buffer was completely removed. Then the protein was freeze-dried for 24 h.

### Protein crystallization

The three different crystal forms, which were also used in our previous backbone-dynamics study although with different isotope labeling (34), were obtained by sitting-drop crystallisation with buffer conditions described below. In all cases, the crystals were obtained using sitting-drop crystallisation plates (Hampton research Cryschem plate, catalog number HR3-159) with a 40 μL sitting drop and 450 μL of reservoir buffer.

For preparing MPD-ub crystals ubiquitin was dissolved in buffer A (20 mM ammonium acetate, pH 4.3) at a concentration of 20 mg/mL. Buffer B (50 mM citrate, pH 4.1) was prepared and mixed with methyl pentanediol (MPD) at a volume ratio of 40:60 (buffer B : MPD), and 450 μL of this mix was placed in the reservoir of the wells. In the sitting drop, 37 μL of the ubiquitin / buffer A solution was mixed with 10 μL of the buffer B / MPD solution. The plate was covered with CrystalClear adhesive tape and kept at 4 ° C. After ca. 1-2 weeks, needle-shaped (“sea-urchin like”) crystals appeared. For preparing cubic-PEG-ub crystals, the reservoir contained 450 μL of buffer C (100 mM 2-(N-morpholino)ethanesulfonic acid (MES), pH 6.3, 20% (weight) PEG 3350 and 100 mM zinc acetate). The protein solution (20 mg/mL of ubiquitin) in buffer D (20 mM ammonium acetate, pH 4.8) was mixed with buffer C at a 1:1 ratio, and 45 μL thereof were placed in the sitting-drop holder. Cubic-shape crystals were obtained within 1 week at ca. 23 ° C.

For preparing rod-PEG-ub crystals, the reservoir buffer contained buffer E (50mM 4-(2-hydroxyethyl)-1-piperazineethanesulfonic acid (HEPES), pH 7.0, 25% PEG 1500 and 25 mM zinc acetate). The protein was dissolved in buffer D at 20 mg/mL, and mixed with reservoir buffer E (1:1), akin to the cubic-PEG-ub procedure.

Protein crystals were transferred to a custom-built ultra-centrifuge tool (essentially a funnel placed on top of a 1.3 mm Bruker NMR rotor, with dimensions that fit the buckets of a Beckman SW32 rotor). The crystals of each kind (ca. 3 mg protein) were centrifuged into their individual rotors, and the rotor caps were glued with two-component epoxy glue to avoid loss of water.

### NMR

All experiments were performed on a Bruker Avance III spectrometer operating at 600 MHz ^1^H Larmor frequency (14.1 T) with a Bruker 1.3 mm probe where the main coil was tuned to ^1^H, ^13^C and ^15^N frequencies, and an auxiliary coil to ^2^H frequency. The MAS frequency was set to 40-50 kHz (specified in the figure panels and below) and maintained constant to within less than 10 Hz. The effective sample temperature was ca. 28 ° C. The temperature was determined from a non-temperature sensitive signal of MPD and the bulk water line, using the equation T[° C] = 455 - 90·δ_H2O_, where δ_H2O_ is the shift of the bulk water line in parts-per-million (ppm).

The pulse sequences for the proton-detected ^1^H-^13^C correlation experiments (hCH) have been presented in Fig. S2 of ref. (28). They include ^1^H excitation, cross-polarisation to ^13^C for chemical-shift editing (with ca. 10 kHz ^1^H WALTZ-16 decoupling), flip-back of ^13^C coherence to ^13^C_z_ for ca. 40 ms water suppression by a train of ^1^H pulses with 18 kHz field-strength amplitude and a duration of 820 μs, alternating in phase (*±* 15 °). The indirect ^13^C dimension was typically sampled for ca. 12-15 ms, using a spectral width of 15 ppm (2250 Hz); the ^1^H dimension was sampled for ca. 50 ms.

The cross-polarisation steps (H to C and C to H) used RF fields of ca. 85 kHz and 35 kHz at 50 kHz MAS frequency, and a duration of 400 μs, with a ramp (90% to 100%) on the ^1^H channel; the specified RF field strength is the value at the mid-point of the ramp. Hard pulses were typically 2.5 to 2.6 μs (^1^H), 3.4 to 3.5 μs (^15^N) and 3.2 μs (^13^C).

The time-shifted REDOR, ^13^C R_1_, ^13^C 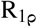 and ^1^H 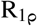 experiments used the same basic hCH correlation experiment, with the appropriate pulse sequence element as described in Fig. S2 of reference (28). In the ^1^H 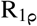 experiment, a spin-lock element was inserted after the initial ^1^H excitation pulse. Except the ^1^H relaxation experiment, all pulse sequences are implemented in NMRlib (61).

The REDOR experiment was performed at a MAS frequency of 44.053 kHz (rotor period of 22.7 μs). For the recoupling pulse train in the REDOR experiment, the ^1^H π and ^13^C π pulses had durations of 5 μs and 6 μs, respectively. The REDOR experiment was implemented with a shift of half of the ^1^H pulses away from the center of the rotor period as described previously (31, 62). The shift of the pulses was such that the shortest time interval between two consecutive ^1^H pulses was 0.5 μs, i.e. the centers of these two consecutive ^1^H pulses were separated by 5.5 μs. Seventeen time points were acquired, from 2 rotor periods to 36 rotor periods in steps of 2 rotor periods (one on each side of the central ^13^C pulse). The REDOR data were collected as a series of 1D spectra.

In the ^13^C 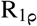 measurements, ten 1D experiments with different spin-lock durations (between 1 and 45 ms) were collected, and this was repeated for 20 different spin-lock RF field strengths, ranging from 2 kHz to 40 kHz. A ^1^H π pulse was applied in the center of the relaxation period to suppress cross-correlated relaxation effects (63). In the ^13^C R_1_ measurements, ten 1D experiments with different relaxation delays (between 1 and 45 ms) were collected. In the ^1^H 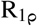 measurements, ten 1D experiments with different spin-lock durations (between 1 and 25 ms) were collected, and this was repeated for 25 different spin-lock RF field strengths, ranging from 1 kHz (for MPD-ub) or 2 kHz (cubic-PEG-ub) to 40 kHz.

To analyse the data, the peaks in the 1D series of spectra were integrated using in-house written python scripts. The relaxation decay profiles were fitted using a simple exponential fit. To interpret the REDOR data, a series of numerical simulations of the REDOR recoupling element was conducted using the GAMMA simulation package (64). The simulations were performed with different values of the tensor anisotropy and biaxiality, resulting in a “2D grid” of simulated time traces. The experimental data were fitted by first calculating Δ *S/S*_ref_ = (*S*_ref_ − *S*_rec_)*/S*_ref_ for both the experiment and the simulation, where *S*_rec_ and *S*_ref_ are the signal intensities in the recoupling experiment and reference experiment (with and without ^1^H pulses, respectively). Then, the experimental curve was compared to each simulation, by calculating a chi-square value as the sum of the squared deviations between the experimental and simulated Δ*S/S*_ref_ divided by the squared experimental error estimate. The best-fit values of the two fitted parameters, tensor anisotropy and biaxiality, were taken to be those for which the calculated chi-square value was minimal over the simulated 2D grid. In an alternative fitting approach, we have performed the simulations with an explicit jump model, assuming that ring flips cause a change of 120° in orientation of ^1^H-^13^C vector. A 1D grid of these simulations was compiled, where the dipolar tensor anisotropy was varied.

Confidence intervals were obtained from Monte-Carlo simulations: the best-fit curve (relaxation decay or REDOR curve) along with experimental uncertainties (of the intensities or Δ *S/S*_ref_ values) were used to generate a “noisy” data set, by choosing the points randomly within a normal distribution around the best-fit data point. One thousand such noisy data sets were fitted using the same procedure as described above, and the standard deviation of the fitted parameters is reported here.

The calculation of relaxation-rate constants in Fig. 4D,E and G used Redfield-theory-based analytical expressions, as outlined in the Supplementary Note 1. The 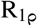 equations have been derived in reference (65). The order parameter used for these calculations was set to *S* = 0.661 (unless stated otherwise), which corresponds to a two-site jump model, *S*^2^ = (3*cos*^2^*ϕ* + 1)*/*4= 0.437, where it is assumed that the jumps occur between the two equiprobable states and cause reorientation of dipolar vector by 120° (66). (Note that 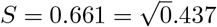 is not exactly the same as *S* = 0.625, which is the scaling of the dipolar-coupling anisotropy from the two-site jumps, used in the REDOR analysis. The latter, however, refers to a different observable and stems from the theoretical description, which also involves a tensor biaxiality.) For all calculations we have used a value of the dipolar coupling that corresponds to 1.09 Å bond length (23327 Hz tensor anisotropy) (16). The CSA tensor was assumed to be uniaxial (axially symmetric), with a value of Δδ =-159 ppm (67).

Note that although the validity of the Redfield theory for slow motions has been debated (22, 68), it appears to be valid for the range of rate constants considered here (69). More specifically, some discrepancies have been detected between Redfield-theory calculations and numerical simulations; however, where the theory appeared to produce incorrect rate constants, it turned out to be due to the fact that decays are multi-exponential. In essence, the deviations were due to misinterpretation of multi-exponential behavior. Caution is required in extracting relaxation rate constants, as discussed in refs. (70, 71).

### MD simulations

Four MD trajectories have been analyzed to glean information on phenylalanine dynamics: those of MPD-ub (2 μs), cubic-PEG-ub (2 μs) and rod-PEG-ub (2 μs) crystals, as well as ubiquitin in solution (8 μs). For example, the MPD-ub trajectory is based on crystallographic structure 3ONS (72) and models the crystalline supercell that is comprised of 4 unit cells. In total, the simulation box contains 24 ubiquitin molecules and 8,772 SPC/E (73) water molecules.

The dimensions of the box were rescaled by a factor 1.016 to reflect the expansion of the crystal at room temperature. For cubic-PEG-ub, the simulated box contains one unit cell, comprising 48 ubiquitin molecules, and for rod-PEG-ub two unit cells, containing the total of 24 ubiquitin molecules in total.

As standard for protein crystal simulations (74), the periodicity of the crystalline lattice is modeled by means of periodic boundary conditions applied to the faces of the simulation box. All crystal trajectories have been recorded in Amber 16 program (75) using ff14SB force field (59); the solution trajectory was recorded in Amber 11 (76) using ff99SB force field (77, 78). Other details of the MD setup can be found in our previous publications (29, 34). MD data have been processed using python library pyxmolpp2 written in-house (available from https://github.com/bionmr-spbu/pyxmolpp2). In particular, this library offers facilities to extract dihedral angles and calculate SASA.

To calculate the chemical shifts, we have used the crystallographic structures 3ONS (72), 3N30 (79) and 3EHV (80) and built the respective crystal lattices. From these lattices we carved out the fragments representing a Ub chain of interest together with the proximal chains. These fragments were subsequently fed into the chemical shift predictor SHIFTX2 (49) to predict the phenylalanine side-chain ^1^H^ε^,^13^C^ε^ chemical shifts. The obtained shifts were averaged over the ε1, ε2 pairs of atoms. In addition, we have also used the high-accuracy NMR structure 1D3Z to similarly predict the chemical shifts in solution.

To calculate the number of flips in the MD simulations, we have employed the following scheme. We defined the flip as the transition between the two states, χ_2_=[60÷180°] and χ_2_=[240÷360°] (cf. Fig. 5). The very rare appearances of the rings in between of these two corridors have been ignored. In this manner we have counted all Phe4 flips observed in all of the crystal and solution simulations and similarly counted all Phe45 flips. These calculations indicate that the flip rate of Phe45 is, on average, 47 times slower than the flip rate of Phe4 in our MD simulations.

## Acknowledgements

The NMR platform in Grenoble is part of the Grenoble Instruct-ERIC center (ISBG; UAR 3518 CNRS-CEA-UGA-EMBL) within the Grenoble Partnership for Structural Biology (PSB), supported by FRISBI (ANR-10-INBS-0005-02) and GRAL, financed within the University Grenoble Alpes graduate school (Ecoles Universitaires de Recherche) CBH-EUR-GS (ANR-17-EURE-0003). This work was supported by the European Research Council (StG-2012-311318-ProtDyn2Function to P.S.) and used the platforms of the Grenoble Instruct Center (ISBG; UMS 3518 CNRS-CEA-UJF-EMBL) with support from FRISBI (ANR-10-INSB-05-02) and GRAL (ANR-10-LABX-49-01) within the Grenoble Partnership for Structural Biology (PSB). We would like to thank Sergei Izmailov for developing and maintaining the pyxmolpp2 library. N.R.S. acknowledges support from St. Petersburg State University in a form of the grant 92425251 and the access to the MRR, MCT and CAMR resource centers.P.S. thanks Malcolm Levitt for pointing out the fact that “tensor asymmetry” is better called “tensor biaxiality”.

All authors would like to make a call for universal peace.

## Author contributions

D.F.G. performed MAS NMR experiments and data analysis and prepared figures. O.O.L. analyzed the MD data, performed chemical shift calculations and prepared figures. N.R.S. participated in the analyses of the MD and chemical shift data, and in writing the manuscript. L.M.B. performed calculations of relaxation rate constants and data fits. I.A. produced the ubiquitin sample. R.L. synthesized the ketoacid precursor. P.S. performed and analyzed NMR experiments, prepared figures and wrote the manuscript with input from all co-authors.

