## Supplementary Information for "Aromatic ring flips in differently packed ubiquitin protein crystals from MAS NMR and MD"

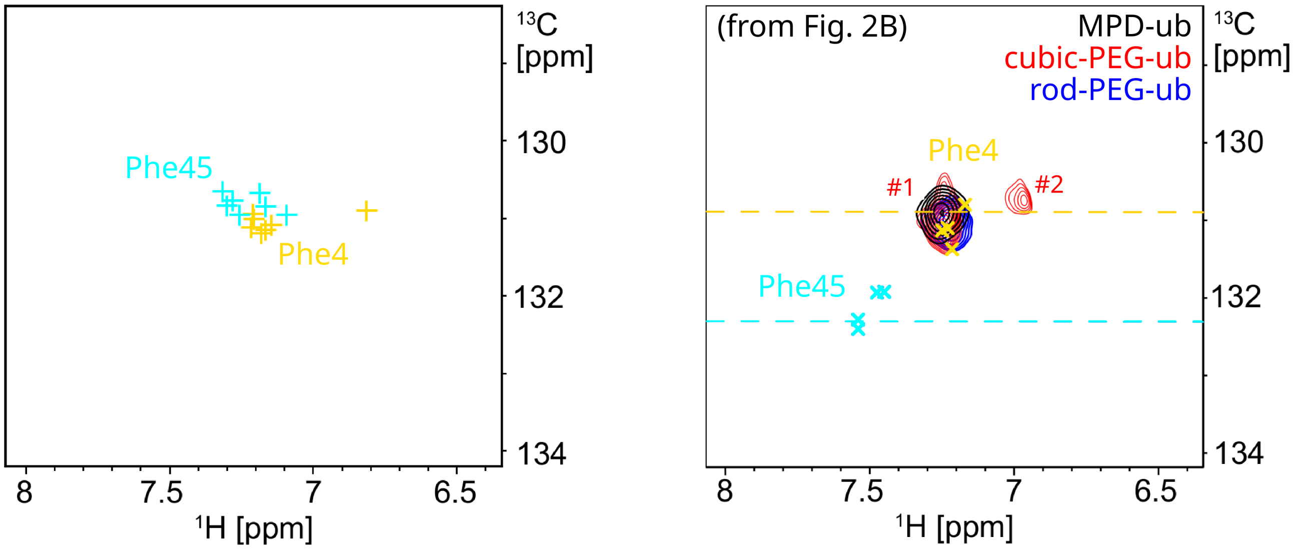


**Fig. S1.** Chemical-shift predictions for $^1H^{\epsilon}$-$^{13}C^{\epsilon}$ spin pairs in Phe4 (colored yellow) and Phe45 (colored cyan) using the program SHIFTX2. It appears that SHIFTX2 correctly predicts the similarity of Phe45 chemical shifts in different crystal forms and in solution. Furthermore, it correctly predicts the Phe4 shifts, including the resonance from chain A in the cubic-PEG-ub crystal, which is shifted due to the intermolecular stacking of Phe4 rings (see Fig.2C). However, SHIFTX2 apparently fails to accurately predict the absolute value of Phe45 shifts as can be deduced from the experimental evidence, see Fig.2B.

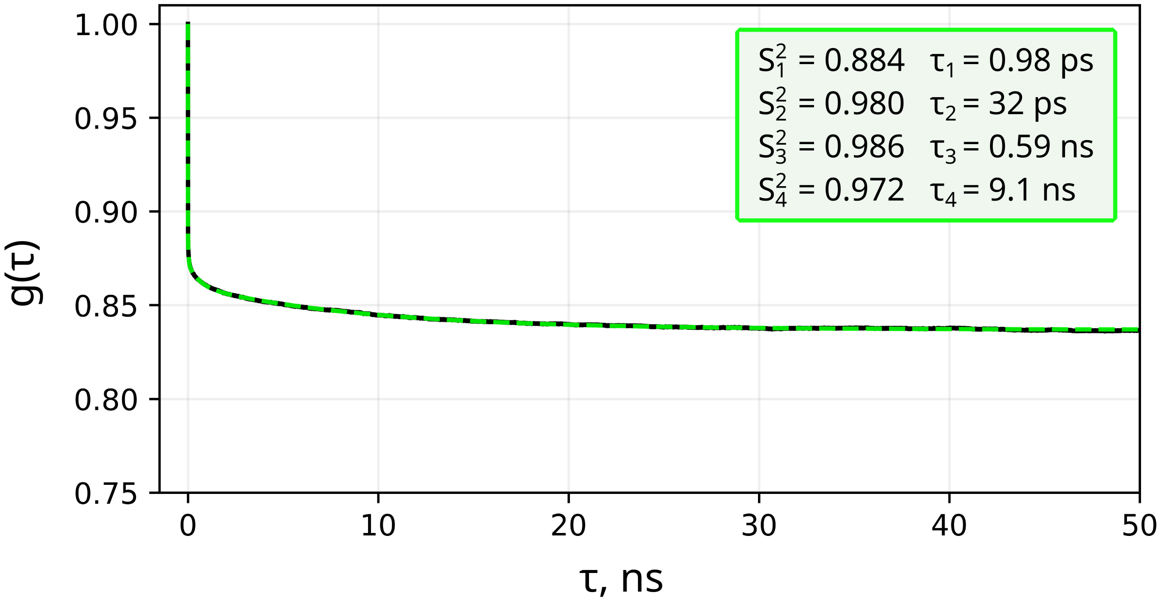


**Fig. S2.** $^{1}H^{\varepsilon}$-$^{13}C^{\varepsilon}$ dipolar correlation function reflecting phenylalanine motions other than ring flips. The data are from six Phe4 residues that do not experience any flips in our MPD-ub simulation (see Fig. 5A). The time correlation function $P_2(\cos\theta)$ (black curve) has been fitted using the 4-exponential decay function parameterized in the spirit of model-free model, Eq. S4 (dashed green curve). The best-fit order parameters and correlation times are listed in the inset. The effect of these non-flip modes on our analyses of spin relaxation rates is illustrated in Fig. S3.

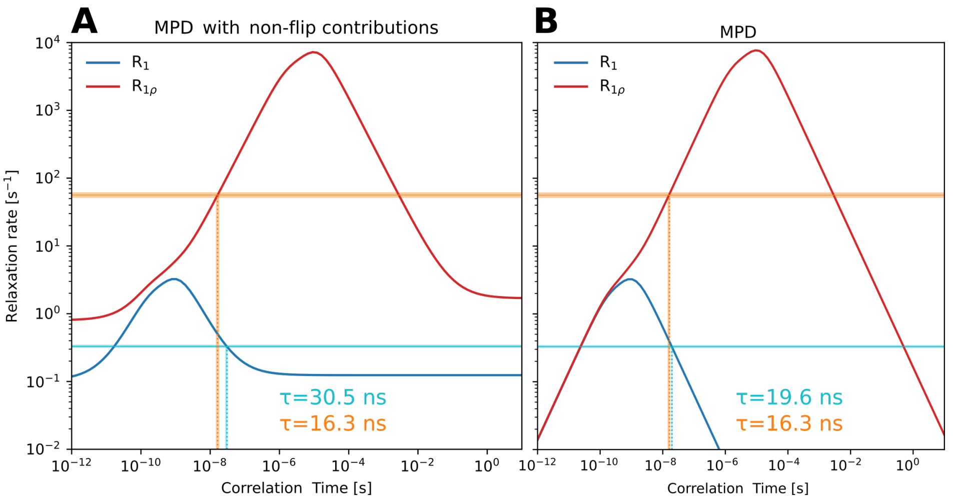


**Fig. S3.** Investigation of the contributions of non-flip motions to the $^{13}C$ relaxation rate constants and their effect on the extracted ring-flip correlation times. (A) Re-analysis of $^{13}C$ $R_{1\rho}$ and $^{13}C$ $R_1$ data from the MPD-ub crystals, which additionally accounts for non-flip motions as listed in Fig. S2. The two profiles in the plot are calculated using Eqns. S8 - S10, see Supplementary Note 2. We used the experimentally observed relaxation rate constants to re-determine the ring-flip time constant $\tau_{flip}$. (B) The original analysis of $^{13}C$ $R_{1\rho}$ and $^{13}C$ $R_1$ data from the MPD-ub crystals, limited to ring-flip dynamics (reproduced from Fig. 4 E). Comparing the results in panels (A) and (B) suggests that taking into consideration non-flip dynamics causes only relatively small change in the determined ring-flip rates.

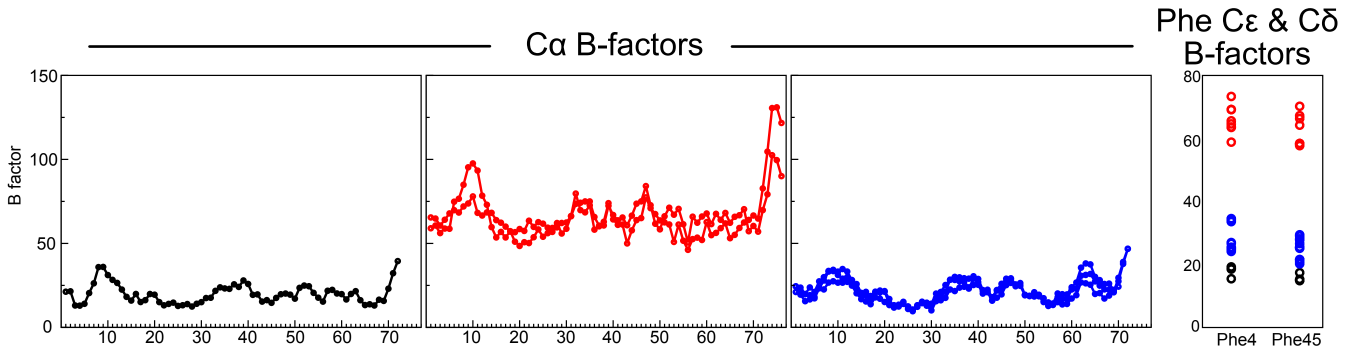


**Fig. S4.** B factors from the crystal structures of MPD-ub (black, PDB 3ONS), cubic-PEG-ub (red, PDB 3N30) and rod-PEG-ub (blue, PDB 3EHV). The left three panels show the $C^{\alpha}$ B factors, and the right panel shows those of the two $C^{\delta}$ and $C^{\varepsilon}$ sites of the Phe rings. Note the large offset of the B factors of cubic-PEG-ub, which we ascribe to overall rocking motion of the protein in the crystal as investigated in ref. (34).

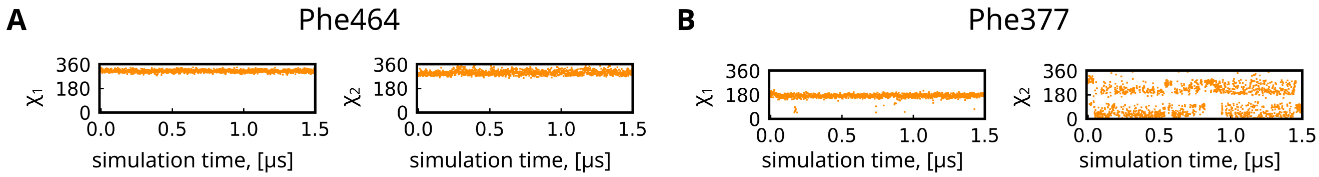


**Fig. S5.** Time traces of side-chain torsional angles $\chi 1$ and $\chi 2$ for surface-exposed residues (A) Phe464 and (B) Phe377 from MD simulation of receptor-binding domain from SARS-CoV-2 spike protein in complex with the mini-protein LCB1 (PDB ID: 7JZU (81)). The 1.5-μs trajectory of this complex was recorded in Amber ff14SB force field using TIP4P-Ew water. Residue Phe464 is in the same rotameric state as Phe4 in ubiquitin, m-85° according to Lovell et al. (58); it is also highly solvent-exposed as evidenced by SASA of 72 Å$^2$ (calculated from the MD trajectory). However, unlike Phe4 in ubiquitin, residue Phe464 shows no flips in our simulations, $\tau_{div} \geqslant 1500$ ns. At the same time, residue Phe377 is in the same rotameric state as Phe45 in ubiquitin, t80°, and solvated at about the same level as Phe45, with SASA of 57 Å$^2$. Yet, unlike Phe45 in ubiquitin, residue Phe377 engages in frequent flips, $\tau_{div} = 10$ ns. These two examples demonstrate that the rotameric state of phenylalanine side chain is not a unique determinant of the flip rate.

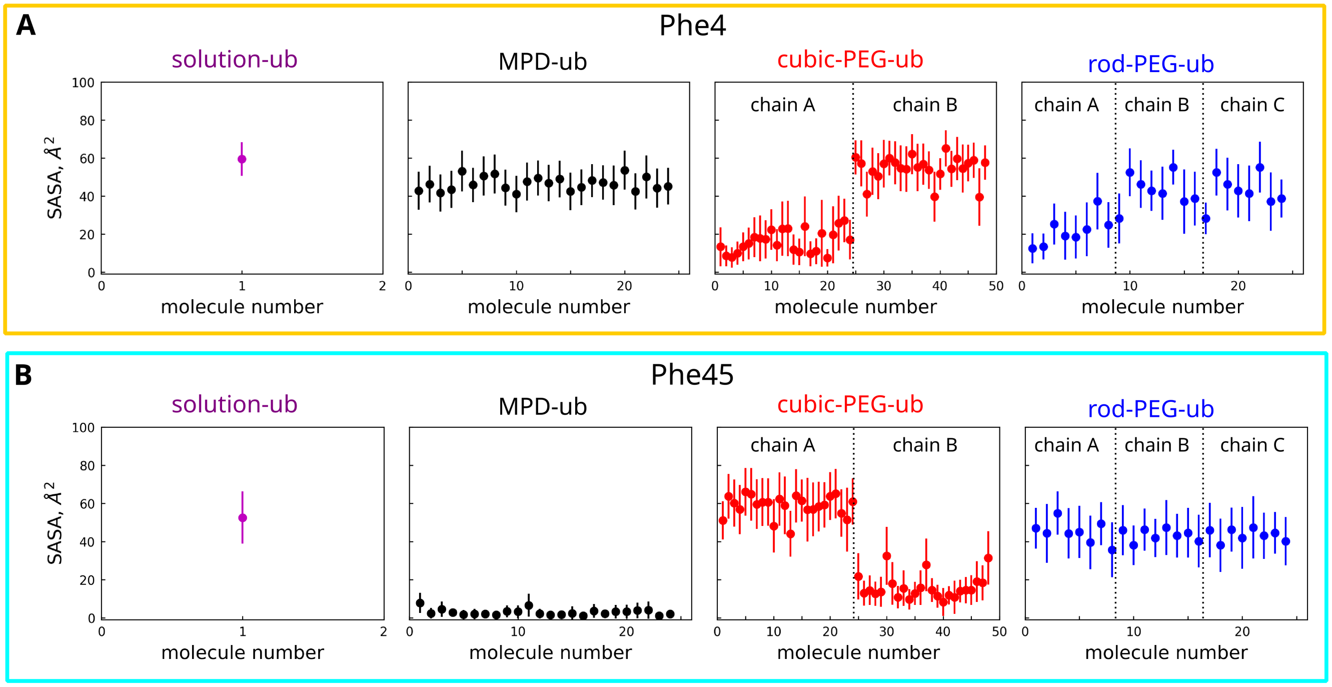


**Fig. S6.** The solvent-accessible surface areas (SASA) of the two phenylalanine residues averaged over 2-μs MD simulations of the three investigated ubiquitin crystals and 8-μs MD simulation of ubiquitin in solution. Of note, the SASA values for Phe4 and Phe45 do not show any significant correlation with the flip rates for these two residues (see Tab. S1). The color coding is the same as elsewhere: black (MPD-ub), red (cubic-PEG-ub), blue (rod-PEG-ub) and magenta (solution form). The MPD-ub simulation box contains 24 ubiquitin molecules that are all classified as chain A. The cubic-PEG-ub simulation box contains 48 ubiquitin molecules equally divided between chain A (molecules 1-24) and chain B (molecules 25-48). The rod-PEG-ub simulation box contains 24 ubiqutin molecules equally divided between chain A (molecules 1-8), chain B (molecules 9-16) and chain C (molecules 17-24).

**Table S1.** Measured $^{13}C^{\varepsilon}$ $R_1$ and $R_{1\rho}$ rate constants for Phe4 in different crystal forms of ubiquitin and the corresponding calculated correlation times $\tau$. Note that the errors on the correlation times are not symmetric, because variance of $R_{1\rho}$ towards higher and lower values propagates to errors in $\tau$ differently for higher and lower $\tau$ values. The $R_{1\rho}$ rate constants are from the measurements at an RF field strength of 30 kHz.

| Crystal | $R_1$ [s$^{-1}$] | $\tau_{R_1}$ [ns] | $R_{1\rho}$ [s$^{-1}$] | $\tau_{R_{1\rho}}$ [ns] |
|---|---|---|---|---|
| **MPD-ub** | 0.33 ±0.01 | $18.7^{+1.4}_{-0.2}$ | 56.5 ±4.1 | $16.3^{+1.2}_{-2.2}$ |
| **cubic-PEG-ub, peak 1** | 0.47 ±0.05 | $13.2^{+2.0}_{-0.9}$ | 53.5 ±5.1 | $15.2^{+1.1}_{-2.0}$ |
| **cubic-PEG-ub, peak 2** | 0.61 ±0.05 | $10.7^{+0.8}_{-0.8}$ | 65.8 ±6.2 | $18.7^{+1.4}_{-2.5}$ |
| **rod-PEG-ub** | 0.55 ±0.03 | $11.5^{+0.9}_{-0.8}$ | 32.5 ±0.7 | $10.0^{+0.1}_{-0.2}$ |

**Table S2.** The characteristic times of Phe flips in ubiqutin according to our MD simulations. The time $\tau_{div}$ is the length of the trajectory divided by the number of flips. The time $\tau_{corr}$ is the flip correlation time, which has been calculated along the lines of Redfield theory treatment. Specifically, we have extracted the time points of all Phe flips from a given MD trajectory and constructed a pseudo-trajectory where dipolar vector (e.g. $H^{\varepsilon}$-$C^{\varepsilon}$) changes its orientation by 120° with each flip. The obtained pseudo-trajectory reproduces Phe ring flips, while eliminating other motional modes such as fast axial fluctuations. The pseudo-trajectory was used to compute the correlation function $P_2(\cos(\theta))$, which was subsequently fitted with $g(\tau) = S^2 + (1 - S^2)exp(-\tau/\tau_{corr})$, where the order parameter is $S^2 = 0.437$, according to the relevant 2-site jump model (66).

| | Phe4 | | Phe45* | |
|---|---|---|---|---|
| | $\tau_{div}(ns)$ | $\tau_{corr}(ns)$ | $\tau_{div}(ns)$ | $\tau_{corr}(ns)$ |
| **solution** | 98 | 66 | 4000 | - |
| **MPD-ub** | 906 | 762 | 48000 | - |
| **cubic-PEG-ub chain A** | 215 | 170 | 4000 | - |
| **cubic-PEG-ub chain B** | 118 | 174 | >48000 | - |
| **rod-PEG-ub chain A** | 236 | 217 | 4000 | - |
| **rod-PEG-ub chain B** | 140 | 242 | >16000 | - |
| **rod-PEG-ub chain C** | 334 | 245 | 8000 | - |

* Because of poor statistics (very few flips), $\tau_{div}$ values for Phe45 are loaded with a large uncertainty, while $\tau_{corr}$ cannot be meaningfully determined.

### Supplementary Note 1: Calculation of proton $R_{1\rho}$ relaxation

The $^1H$ $R_{1\rho}$ relaxation can be described by a sum of heteronuclear dipolar $^1H$-$^{13}C$ ($R_{1\rho}{}^{HC}$), homonuclear dipolar $^1H$-$^1H$ ($R_{1\rho}{}^{HH}$) and the $^1H$ CSA ($R_{1\rho}{}^{CSA}$) contributions (82):

$$R_{1\rho}^{\mathrm{HC}} = \frac{1}{4} d_{\mathrm{HC}}{}^2 \left\{ \frac{1}{3}\sin^2(\beta_{\mathrm{eff}}) \Big[ J(\omega_{\mathrm{e}} - 2\omega_{\mathrm{r}}) + 2J(\omega_{\mathrm{e}} - \omega_{\mathrm{r}}) + 2J(\omega_{\mathrm{e}} + \omega_{\mathrm{r}}) + J(\omega_{\mathrm{e}} + 2\omega_{\mathrm{r}}) + 9J(\omega_{\mathrm{C}}) \Big] + \frac{1}{4}(3 + \cos(2\beta_{\mathrm{eff}})) \Big[ 3J(\omega_{\mathrm{H}}) + 6J(\omega_{\mathrm{H}} + \omega_{\mathrm{C}}) + J(\omega_{\mathrm{H}} - \omega_{\mathrm{C}}) \Big] \right\} \quad \textbf{(S1)}$$

$$R_{1\rho}^{\mathrm{HH}} = \frac{3}{4} d_{\mathrm{HH}}{}^2 \left\{ \frac{1}{8}\sin^2(2\beta_{\mathrm{eff}}) \Big[ J(\omega_{\mathrm{r}} - \omega_{\mathrm{e}}) + J(\omega_{\mathrm{r}} + \omega_{\mathrm{e}}) \Big] + \frac{1}{2}\sin^4(\beta_{\mathrm{eff}}) \Big[ J(\omega_{\mathrm{r}} - 2\omega_{\mathrm{e}}) + J(\omega_{\mathrm{r}} + 2\omega_{\mathrm{e}}) \Big] + \frac{1}{16}\sin^2(2\beta_{\mathrm{eff}}) \Big[ J(2\omega_{\mathrm{r}} - \omega_{\mathrm{e}}) + J(2\omega_{\mathrm{r}} + \omega_{\mathrm{e}}) \Big] + \frac{1}{4}\sin^4(\beta_{\mathrm{eff}}) \Big[ J(2\omega_{\mathrm{r}} - 2\omega_{\mathrm{e}}) + J(2\omega_{\mathrm{r}} + 2\omega_{\mathrm{e}}) \Big] + \frac{1}{4}(7 - 3\cos(2\beta_{\mathrm{eff}})) \Big[ J(\omega_{\mathrm{H}}) \Big] + \frac{1}{2}(5 + 3\cos(2\beta_{\mathrm{eff}})) \Big[ J(2\omega_{\mathrm{H}}) \Big] \right\} \quad \textbf{(S2)}$$

$$R_{1\rho}^{\mathrm{CSA}} = \frac{1}{3}\omega_{\mathrm{H}}^2 \Delta\sigma^2 \left\{ \frac{1}{9}\sin^2(\beta_{\mathrm{eff}}) \Big[ J(\omega_{\mathrm{e}} - 2\omega_{\mathrm{r}}) + 2J(\omega_{\mathrm{e}} - \omega_{\mathrm{r}}) + 2J(\omega_{\mathrm{e}} + \omega_{\mathrm{r}}) + J(\omega_{\mathrm{e}} + 2\omega_{\mathrm{r}}) \Big] + \frac{1}{4}(3 + \cos(2\beta_{\mathrm{eff}}) \Big[ J(\omega_{\mathrm{H}}) \Big] \right\} \quad \textbf{(S3)}$$

Here, $d_{\mathrm{HC}}$ and $d_{\mathrm{HH}}$ are the hetero- and homonuclear dipolar couplings given by $d_{12} = -\frac{\mu_0}{4\pi}\frac{\gamma_1\gamma_2\hbar}{r_{12}^3}$ with the vacuum permeability $\mu_0$, the gyromagnetic ratios $\gamma_i$, Planck's constant $\hbar$ and the internuclear distance $r_{12}$. $\Delta\delta = \frac{3}{2}(\delta_{zz} - \delta_{iso})$ is the reduced chemical shift anisotropy, $\omega_{\mathrm{e}}$ is the amplitude of the effective field, $\beta_{\mathrm{eff}}$ is the tilt angle of the effective field, $\omega_{\mathrm{r}}$ is the MAS frequency, and $\omega_{\mathrm{H}}$ and $\omega_{\mathrm{C}}$ are the $^1H$ and $^{13}C$ Larmor frequencies, respectively. The spectral densities $J(\omega)$ are defined as $J(\omega) = \frac{2}{5}(1 - S^2)\frac{\tau_{\mathrm{c}}}{1 + (\omega\tau_{\mathrm{c}})^2}$, with the order parameter $S$ and the correlation time $\tau_{\mathrm{c}}$.

For the homonuclear dipolar relaxation of an $H^{\varepsilon}$ proton (Fig. 4G), we considered a four-spin system inspired by the environment of Phe45 in the current deuterated, selectively $^1H$,$^{13}C$-labeled sample. This system contains the $H^{\varepsilon}$ proton of interest, its directly bound $C^{\varepsilon}$, another $H^{\varepsilon}$ proton and the closest $H^N$ amide proton in the backbone (Thr66). The distances between the nuclei are 1.09 Å for $C^{\varepsilon}$-$H^{\varepsilon}$, 4.3 Å for $H^{\varepsilon}$-$H^{\varepsilon}$ and 3.0 Å for $C^{\varepsilon}$-$H^N$. In addition to the dipolar couplings that result from these spin pairs, we have taken into account the $H^{\varepsilon}$ CSA. The proton chemical shift anisotropy $\Delta\delta$ was set to 9 ppm (83). The relaxation rates were calculated assuming a magnetic field corresponding to 600 MHz proton Larmor frequency, 44.053 kHz MAS, an effective field of 80 kHz and an effective tilt angle of 90°. The order parameters $S$ were set to 0.661 for the heteronuclear $^{13}C^{\varepsilon}$-$^1H^{\varepsilon}$ relaxation, 1.0 for the homonuclear $^1H^{\varepsilon}$-$^1H^{\varepsilon}$ relaxation and 0.5 for the homonuclear $^1H^{\varepsilon}$-$^1H^N$ relaxation (66).

### Supplementary Note 2: Including the effect of motional modes other than ring flips

In addition to ring flips, $^{13}C^{\varepsilon}$ spin relaxation is also sensitive to other motional modes, such as small-amplitude axial fluctuations involving $\chi_1$ and $\chi_2$. To estimate the effect of these other modes on the outcome of our analyses, Fig. 4E, we have applied the following scheme. In MPD-ub trajectory, we have identified 6 ubiquitin molecules where Phe4 ring does not experience any flips. For these six sites, we have calculated the average $^1H^{\varepsilon}$-$^{13}C^{\varepsilon}$ dipolar correlation function $g(\tau)$, shown as black curve in Fig. S2. The curve has a familiar shape, decaying toward the plateau value of 0.84; similar shapes are found for backbone sites, where dynamics is also limited to small-amplitude fluctuations (84). As is generally the case, the decay of $g(\tau)$ has a multi-exponential character (85). Accordingly, the function can be fitted with a combination of several exponentials in the spirit of the extended Lipari-Szabo model (86). Here we have found that four exponential terms are necessary and sufficient to fit $g(\tau)$. For the purpose of our analyses, we have assumed that there are four independent motional modes characterized by their respective order parameters $S_i^2$ and correlation times $\tau_i$. We have further assumed that the net correlation function is a product of the four mode-specific functions (87):

$$g_{fit}(\tau) = ((1 - S_1^2)exp(-\tau/\tau_1) + S_1^2) \cdot ((1 - S_2^2)exp(-\tau/\tau_2) + S_2^2) \cdot ((1 - S_3^2)exp(-\tau/\tau_3) + S_3^2) \cdot ((1 - S_4^2)exp(-\tau/\tau_4) + S_4^2) \quad \textbf{(S4)}$$

The fitting of the correlation function $g(\tau)$ with the ansatz from Eq. S4 is illustrated in Fig. S2 (dashed green curve in the plot). The obtained time constants, $\tau_1$ = 0.98 ps, $\tau_2$ = 32 ps, $\tau_3$ = 0.59 ns, $\tau_4$ = 9.2 ns are different by an order of magnitude, thus confirming the statistical independence of the respective motional modes. The first time scale, $\tau_1$, is extremely short and, in fact, determined by the time interval that is used to record protein coordinates during the MD simulation. This time scale is clearly not relaxation-active and, therefore, the expression fo $g_{fit}(\tau)$ can be simplified to:

$$g_{red}(\tau) = S_1^2 \cdot ((1 - S_2^2)exp(-\tau/\tau_2) + S_2^2) \cdot ((1 - S_3^2)exp(-\tau/\tau_3) + S_3^2) \cdot ((1 - S_4^2)exp(-\tau/\tau_4) + S_4^2) \quad \textbf{(S5)}$$

Furthermore, the term $S_1^2$, which pertains to sub-picosecond vibrations and librations, must be already factored into the interaction constants (dipolar and CSA) used in the data analyses. Therefore, the relevant portion of the correlation function is reduced to:

$$g_{red}(\tau) = ((1 - S_2^2)exp(-\tau/\tau_2) + S_2^2) \cdot ((1 - S_3^2)exp(-\tau/\tau_3) + S_3^2) \cdot ((1 - S_4^2)exp(-\tau/\tau_4) + S_4^2) \quad \textbf{(S6)}$$

Using this MD-derived correlation function, we now construct a more general correlation function, which additionally accounts for the effect of ring flips:

$$G(\tau) = ((1 - S_2^2)exp(-\tau/\tau_2) + S_2^2) \cdot ((1 - S_3^2)exp(-\tau/\tau_3) + S_3^2) \cdot ((1 - S_4^2)exp(-\tau/\tau_4) + S_4^2) \cdot ((1 - S_{flip}^2)exp(-\tau/\tau_{flip}) + S_{flip}^2) \quad \textbf{(S7)}$$

In this expression, the flip order parameter is $S_{flip}^2 = 0.437$ (66), the values of $S_2^2, \tau_2$, $S_3^2, \tau_3$ and $S_4^2, \tau_4$ are fixed according to the MD fitting results (Fig. S2), and $\tau_{flip}$ is the sole variable. It is straightforward to convert Eq. S7 into formula for spectral density $J(\omega)$, but the resulting expression is rather bulky (not shown). Alternatively, one can notice that in our case $\tau_2 \ll \tau_3 \ll \tau_4, \tau_{flip}$, which permits the following simplification:

$$J(\omega) = \frac{2}{5}[(1 - S_2^2)\frac{\tau_2}{1 + (\omega\tau_2)^2} + S_2^2(1 - S_3^2)\frac{\tau_3}{1 + (\omega\tau_3)^2} + S_2^2 S_3^2 \xi] \quad \textbf{(S8)}$$

$$\xi = (1 - S_4^2)(1 - S_{flip}^2)\frac{\tau'}{1 + (\omega\tau')^2} + (1 - S_4^2)S_{flip}^2\frac{\tau_4}{1 + (\omega\tau_4)^2} + S_4^2(1 - S_{flip}^2)\frac{\tau_{flip}}{1 + (\omega\tau_{flip})^2} \tag{S9}$$

$$\tau' = (\frac{1}{\tau_4} + \frac{1}{\tau_{flip}})^{-1} \tag{S10}$$

Note that this simplification is valid only for our specific case, where the time scales $\tau_2$, $\tau_3$ and $(\tau_4, \tau_{flip})$ are separated by at least an order of magnitude, see Fig. S2 and Tab. S1.

Eqns. S8 - S10 have been used to re-interpret our experimental results, arriving at Fig. S3. Comparing this latter to Fig. 4E in the main text, one can appreciate the significance of motional modes other than ring flips and quantify the bias in determination of $\tau_{flip}$ due to the neglect of those other motional modes.

Finally, note that MD-derived parameters of local dynamics are not always accurate; the specific set of parameters used in our calculations pertain to one specific crystal form, that of MPD-ub. Furthermore, different motional modes underlying the correlation function may not be completely independent, but rather may prove to be partially correlated. In addition, the relevant constants, e.g. $r_{CH}$ = 1.09 Å, are also not very accurately known and are a subject of debate focusing on the effect of vibrational averaging (88, 89). Thus, the results in Fig. S3 are, at best, a semi-quantitative estimate.
